# Effect of the LSD1 Inhibitor RN-1 on γ-globin and Global Gene Expression during Erythroid Differentiation in Baboons (*Papio anubis*)

**DOI:** 10.1101/2023.07.28.550962

**Authors:** Vinzon Ibanez, Kestis Vaitkus, Maria Armila Ruiz, Zhengdeng Lei, Mark Mannschein-Cline, Zarema Arbieva, Donald Lavelle

## Abstract

Increased levels of Fetal Hemoglobin reduce the severity of symptoms of patients with sickle cell disease and decrease the risk of death. An affordable, small molecule drug that stimulates HbF expression in vivo would be ideally suited to treat the large numbers of SCD patients that exist worldwide. Our previous work showed that administration of the LSD1 (KDM1A) inhibitor RN-1 to normal baboons increased Fetal Hemoglobin (HbF) and was tolerated over a prolonged treatment period. HbF elevations were associated with changes in epigenetic modifications that included increased levels of H3K4 di-and tri-methyl lysine at the γ-globin promoter. While dramatic effects of the loss of LSD1 on hematopoietic differentiation have been observed in murine LSD1 gene deletion and silencing models, the effect of pharmacological inhibition of LSD1 in vivo on hematopoietic differentiation is unknown. The goal of these experiments was to investigate the in vivo mechanism of action of the LSD1 inhibitor RN-1 by determining its effect on γ-globin expression in highly purified subpopulations of bone marrow erythroid cells enriched for varying stages of erythroid differentiation isolated directly from baboons treated with RN-1 and also by investigating the effect of RN1 on the global transcriptome in a highly purified population of proerythroblasts. Our results show that RN-1 administered to baboons targets an early event during erythroid differentiation responsible for γ-globin repression and increases the expression of a limited number of genes including genes involved in erythroid differentiation such as GATA2, GFi-1B, and LYN.

## Introduction

The switch from Fetal Hemoglobin (HbF) to Adult Hemoglobin (HbA) expression observed during development is recapitulated during adult bone marrow erythroid differentiation [1–3]. During this process, repressive epigenetic marks are acquired and activating epigenetic marks removed from the γ-globin promoter by epigenome-modifying enzymes such as histone deacetylases (HDACs), DNA methyltransferase (DNMT1), LSD1 (KDM1A), and G9A (EHMT2) [4,5]. These enzymes are functional components of multiprotein co-repressors recruited to the γ-globin gene promoter by the trans-acting repressors TR2/TR4, BCL11A, and ZBTB7A [6–11]. Various pharmacological inhibitors targeting HDACs, DNMT1, LSD1, and G9A increase γ-globin expression in different experimental model systems and some of have advanced to clinical trials [12–14]. LSD1 inhibitors are particularly potent HbF activators and promising therapeutic agents as shown by studies in primary erythroid cell cultures derived from normal [15] and β°-thalassemia/HbE patients [16], sickle cell disease mouse models [17–19] and baboons [20–22], but detailed knowledge of their mechanism of action has not been established. Our previous studies showed that the LSD1 inhibitor RN-1 administration stimulated impressive increases in HbF levels [20] and was generally well-tolerated during a prolonged treatment period in baboons. [21] Murine gene deletion and knockdown models have shown dramatic effects of the loss of LSD1 on hematopoietic differentiation. [23–26] To what extent in vivo pharmacological inhibition of LSD1 using doses sufficient to increase HbF to clinically relevant levels in a non-human primate animal model affects gene expression and erythroid differentiation is unknown. Baboons are considered the best animal model to test the ability of drugs to increase HbF due to conservation of developmental regulation and structure of the β-globin gene complex between baboons and man [27–29]. Experimental findings of HbF-inducing drugs in baboons [30, 31] are predictive of responses in man and have allowed direct translation of experimental baboon studies to clinical trials in patients with sickle cell disease [32–37]. The goal of this study was to investigate the in vivo mechanism of action of RN-1 in baboons by determining the effect of RN-1 administration in vivo on γ-globin expression during erythroid differentiation in highly purified subpopulations of bone marrow erythroid cells isolated directly from baboons treated with RN-1 and also its effect on the global transcriptome. Methods to isolate subpopulations of bone marrow (BM) cells highly enriched at different stages of erythroid differentiation from fresh baboon BM aspirates obtained from resting, normal controls and from RN-1 treated baboons were developed using a combination of immunomagnetic column selection and FACS. RT-PCR analysis of γ-globin expression within purified BM erythroid subpopulations showed that expression of γ-globin relative to β-globin mRNA (γ/γ+β) in untreated baboons was 4 fold higher in BFUe than in CFUe and 11 fold higher in BFUe than in proerythroblasts consistent with repression of the γ-globin gene during the BFUe to CFUe transition. RN-1 treatment increased γ-globin expression (γ/γ+β) 4 fold in BFUe, 25 fold in CFUe, and >100 fold in proerythroblasts compared to untreated controls, indicating that RN-1 interfered with initiation of repression of the γ-globin gene at the early stages of differentiation. RNAseq analysis of FACS-purified BM proerythroblasts of untreated control (n=2) and RN-1 treated baboons (0.25mg/kg/d; 3d; n=4; 2 treated for 3d; 2 baboons treated >265d) showed that the expression of a limited number genes was increased in both the short-term (3d) and long-term (>265d) treated baboons and within this subset were genes that have major effects on erythroid differentiation such as, GATA2, GFi1B, and LYN. Additional flow cytometry analysis of baboon BM aspirates obtained from RN-1-treated and untreated baboons showed that the drug increased the proportion of proerythroblasts and decreased the proportion of terminal erythroid BM cells. We conclude that the LSD1 inhibitor RN-1 administered to baboons at doses that increases HbF to clinically significant levels 1) targets an early event responsible for γ-globin repression to increase γ-globin expression, and 2) increases the expression of a limited set of genes including, GATA2, GFi-1B, and LYN associated with an inhibition of erythroid differentiation.

## Materials and Methods

### Ethics Statement

All procedures involving baboons were approved by the Animal Care Committee of the University of Illinois at Chicago (ACC #20-152).

### Baboons

Baboons were housed at the University of Illinois at Chicago Biologic Resources Laboratory (UIC BRL) under conditions that meet the Association for Assessment and Accreditation of Laboratory Animal Care (AAALAC) standards. The University of Illinois at Chicago is committed to the judicious, humane use of animals in research and teaching and is also committed to maintaining full accreditation by the AAALAC. In accordance with this commitment, all measures are taken to socially house nonhuman primates in pairs or groups in accordance with Section 3.81(a) of the Animal Welfare Standards, when stable pairs can be established or unless otherwise exempt due to veterinary or scientific reasons in facilities that meet AAALAC standards for accreditation. Nonhuman primates receive a variety of foodstuffs in addition to their daily diet of primate biscuits which may include seasonal fruits and vegetables, Softies, and human food treats. They also receive a foraging mixture consisting of a variety of seeds, dried fruit, and dry dog food several times each week, and several times each month receive a novel food enrichment item which may include seasonal fruits (green onions, radishes, melons, oranges, holiday candies, etc.) and treats (frozen juice popsicles popcorn balls etc.). An adequate water supply is continuously available. RN-1 (Cayman Chemical Company) was administered to baboons (0.25mg/kg/d for three consecutive days and from baboons treated with RN-1 (0.25mg/kg/d; 5d/wk) long-term (>265d) by subcutaneous injection. Bone marrow aspirates were obtained from both normal resting baboons and RN-1-treated baboons 48 hours following the last day of drug administration. Bone marrow aspirations were performed from the hips of animals under ketamine/xylazine anesthesia (10mg/kg; 1mg/kg). Prior to bone marrow sampling, buprenex (0.01mg/kg IM) was given to alleviate potential pain and suffering. All baboon studies and procedures were approved by the Animal Care Committee of the University of Illinois at Chicago (ACC# 20-152).

### Purification and analysis of bone marrow subpopulations

Mononuclear cells were isolated from baboon bone marrow aspirates following sedimentation on Percoll gradients. Cells were incubated with CD105+ (Invitrogen; clone SN6) for 20 minutes at 4°C followed by three washes in phosphate-buffered saline (PBS). The cell pellets were then incubated with immunomagnetic-conjugated rat anti-mouse IgG microbeads (Miltenyi) and the CD105+ cells isolated using an LS immunomagnetic column. The isolated CD105+ cells were then incubated with PE-conjugated mouse anti-human CD34 (BD Bioscience; clone 563), FITC-conjugated mouse anti-human CD117 (BD Bioscience; clone 104D2) and APC-conjugated mouse-anti-baboon red blood cell antigen (BD Bioscience, clone E34-731). Subpopulations of cells were isolated by FACS-purification using a MoFlo Astrios cell sorter at the University of Illinois at Chicago Research Resources Center. Analysis of BM erythroid differentiation by flow cytometry was performed using a Beckman-Coulter FC500 instrument. Colony forming assays were performed by plating 200 cells of each isolated subpopulation in Complete Methylcellulose Media (R&D Systems; HSC003).

### Globin gene expression

RNA was isolated using the RNeasy Minikit (Qiagen #74104) and treated with RNase-free DNase I (Ambion #AM1906) according to the manufacturer’s instructions. Purified RNA was used for the synthesis of cDNA using RevertAid First Strand CDNA Synthesis Kits (Thermo Scientific #K1622). Real time PCR analysis was conducted using custom designed primer-probe combinations were used for analysis of globin transcripts (S1 Table). Absolute numbers of globin transcripts were determined by extrapolation from standard curves prepared from the cloned amplicons [38].

### Global gene expression

Global gene expression analysis was performed by the Genome Research Core at the Research Resources Center of the University of Illinois at Chicago. RNA was isolated from the baboon BM CD105+CD34-CD117+bRBC+ FACS-purified subpopulation (proerythroblasts) using Ambion kits. Purified RNA was provided to the UIC-Genome Research Core for RNAseq analysis. Sequencing libraries were prepared using strand-specific QuantSeq 3’ RNAseq kit (Lexogen, FWD, Cat. No. 015). The QuantSeq protocol generates only one fragment per transcript, resulting in accurate gene expression values, and the sequences obtained are close to the 3’ end of the transcripts. Total RNA in the amount of 50 nanograms per sample was used as an input. Library construction was performed according to the manufacturer’s protocol. In brief, during the first strand cDNA synthesis, an oligo dT primer containing an Illumina-compatible sequence at its 5’ end was hybridized to mRNA, and reverse transcription was performed. After that, the RNA template was degraded, and during second strand synthesis, the library was converted to dsDNA. Second-strand synthesis was initiated by a random primer containing an Illumina-compatible linker sequence at its 5’ end. The double-stranded libraries were purified by using magnetic beads to remove all reaction components. Next, the libraries were amplified to add the complete adapter sequences required for cluster generation and to generate sufficient material for quality control and sequencing. A number of PCR amplification cycles was 18, as determined by test qPCR using a small pre-amplification library aliquot for each individual sample. Final amplified libraries were purified, quantified, and fragment sizes were confirmed to be within 275-300 bp by gel electrophoresis using Agilent 4200 TapeStation (D5000 Screen Tape). Concentration of the final library pool was confirmed by qPCR. Sequencing was performed on NextSeq 500 (Illumina), high-output kit, 1×75 nt single-end reads.

Raw reads were aligned to reference genome Panu_2.0 using BWA MEM [39]. Gene expression was quantified using FeatureCounts [40]. Differential expression statistics (fold-change and *p*-value) were computed using edgeR, [41,42] including both multi-group comparisons using generalized linear models as well as pair-wise tests between experimental groups using the exactTest function. In all cases we adjusted *p*-values for multiple testing using the false discovery rate (FDR) correction of Benjamini and Hochberg [43].

Sequence data has been deposited in the GEO database under the GEO accession number GSE235633.

### Statistical Analysis

Mean, standard deviation and *p*-values for colony assays and RT-PCR assays were calculated using the Student T test two-tailed for unpaired samples using Graphpad software.

## Results

### Isolation of baboon bone marrow erythroid cells

Bone marrow aspirates from normal baboons were separated by sedimentation using Percoll gradients followed by positive selection of CD105+ cells using immunomagnetic antibody columns. The CD105+ population was then further fractionated into six subpopulations based upon expression of CD34, CD117, and the baboon red blood cell antigen (bRBC) that were designated R6 (CD34+CD117+bRBC-), R8 (CD34+CD117-bRBC-), R10 (CD34-CD117+bRBC-), R5 (CD34+CD117+bRBC+), R7 (CD34-CD117+bRBC+), and R9 (CD34-CD117-bRBC+) (Fig 1A). Morphological analysis of Wright’s-stained cytospin preparations and methylcellulose colony forming assays showed that R8 (CD34+CD117-bRBC-) subpopulation consisted of nonerythroid progenitors but that other the subpopulations were highly enriched for erythroid cells representative of specific stages of erythroid differentiation (Fig 1B). The R7 (CD34-CD117+bRBC+) subpopulation consisted of primarily proerythroblasts while the R9 (CD34-CD117-bRBC+) subpopulation was enriched in orthochromatic and polychromatic erythroid precursors. Colony assays performed to assess the distribution of BFUe and CFUe within the subpopulations (Fig 1C and D) showed that the most primitive subpopulation, R6 (CD34+CD117+bRBC –), contained high numbers of BFUe (69.8±10.8 colonies/200 cells plated; mean±SD) and lesser numbers of CFUe (28.5±14.1 colonies/200 cells plated; mean±SD) with 2.4 X more BFUe than CFUe. Both the R5 (CD34+CD117+bRBC+) and R10 (CD34-CD117+bRBC-) subpopulations were highly enriched in CFUe (R5:66.7±16.3, R10: 71.7±20.2 CFUe/200 cells plated; mean±SD) while the numbers of colony forming cells were very low or absent in the R5

**Fig 1.**
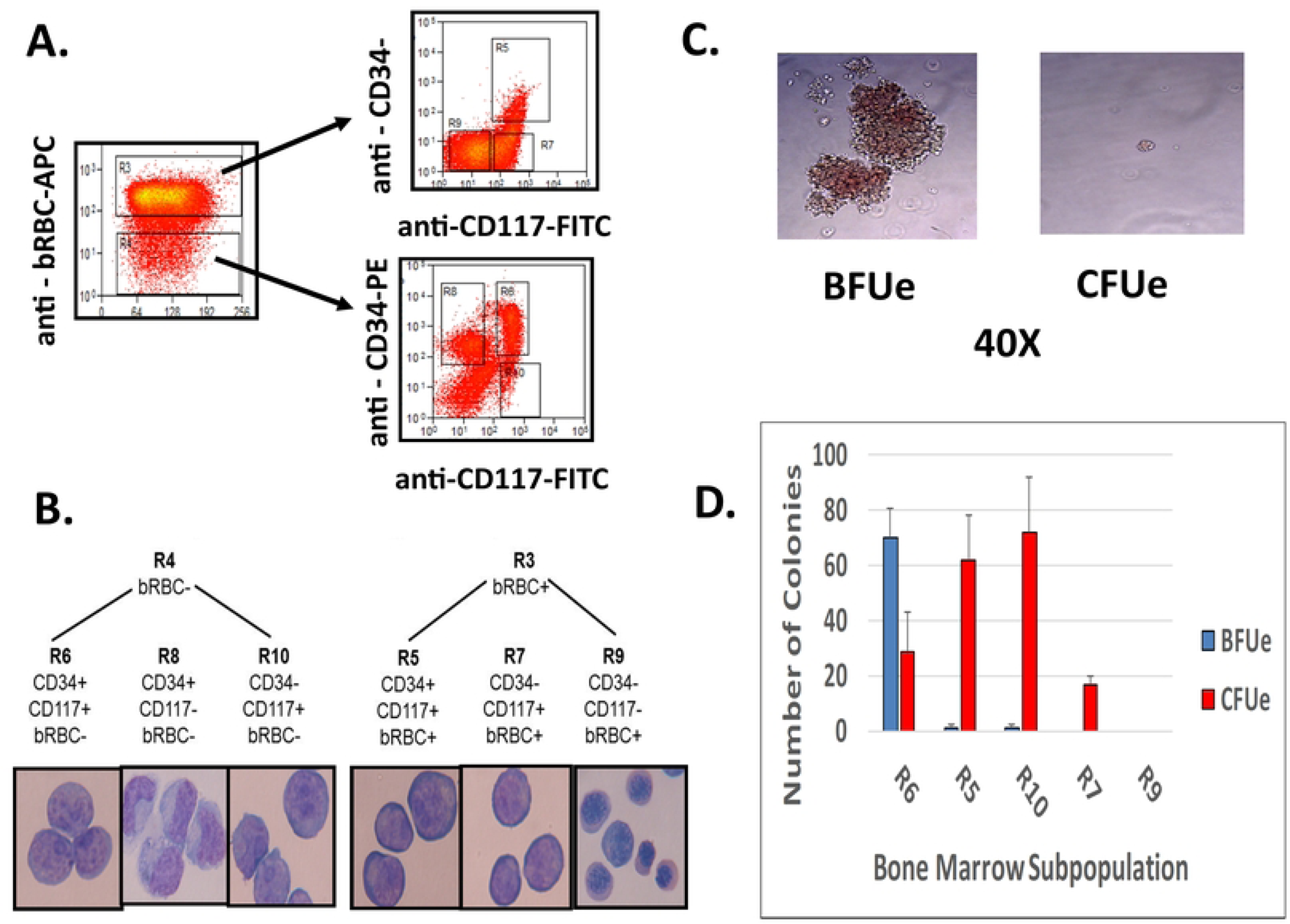
Isolation of Baboon Bone Marrow Erythroid Cell Subpopulations. **A.** Representative FACS plots showing separation of baboon BM erythroid cell subpopulations. **B.** Wright’s stained cytospin preparation of isolated BM erythroid subpopulations. **C.** Morphology of BFUe and CFUe colonies identified in the R6 (CD34+CD117+bRBC-) and R5 (CD34+CD117+bRBC+) subpopulations, respectively shown at 40X magnification. **D.** Results of methylcellulose colony assays showing enrichment of BFUe and CFUe within the various BM subpopulations (200 cells plated; mean±SD).

(CD34-CD117+bRBC+) and R9 (CD34-CD117-bRBC+) subpopulations.

### Effect of RN-1 on γ-globin expression during erythroid differentiation

Expression of the γ-globin gene relative to the β-globin gene was measured by RT-PCR assays in 4 subpopulations of bone marrow cells enriched for different stages of erythroid differentiation (Fig 2A). These subpopulations included the R6 (CD34+CD117+bRBC-) BFUe-enriched subpopulation, the R5 (CD34+CD117+bRBC+) CFUe enriched subpopulation, the R7 (CD34-CD117+bRBC+) proerythroblast enriched subpopulation, and the R9 (CD34-CD117-bRBC+) orthochromatic and polychromatic cell erythroid cell enriched subpopulation. RT-PCR assays showed the relative level of γ-globin to β-globin transcripts was (γ/γ+β) was 6.7 fold higher in the R6 (CD34+CD117+bRBC-) BFUe-enriched subpopulation (0.040+0.016 γ/γ+β) compared to the R5 (CD34+D117+bRBC+) CFUe-enriched R5 subpopulation (0.006+0.004 γ/γ+β; *p*=0.024) and 13 fold higher than R7 (CD34-CD117+bRBC+;) proerthroblast population (0.003+0.002 γ/γ+β; *p*=0.015; Fig 2A). The relative level of γ-globin to GAPDH transcripts in the less differentiated R6 BFUe-enriched subpopulation was 90 fold less than in the R9 terminal erythroid subpopulation (*p*<0.001; Fig 2B). The high level of γ-globin to β-globin RNA ratio was not sustained in the more differentiated populations and is consistent with repression of the γ-globin gene during the BFUe to CFUe transition. The effect of RN-1 on globin gene expression within these subpopulations was assessed (Fig 2A). Baboons were treated with RN-1 (0.25mg/kg/d; 3d) and bone marrow aspirates were obtained 48 hours following the last administration of RN-1. The BM erythroid cell subpopulations were isolated as described above. The levels of γ-globin relative to β-globin (γ/γ+β) RNA in all subpopulations isolated from RN-1 treated baboons (R6: 0.106+0.029 γ/γ+β; R5: 0.136+0.037 γ/γ+β; R7: 0.176+0.059 γ/γ+β; R9:0.089+0.008 γ/γ+β) were significantly higher than in the corresponding subpopulations isolate from untreated controls (R6: 2.6 fold higher, *p*=0.013; R5: 22.6 fold higher, *p*=0.003; R7: 87.5 fold higher, *p*<0.001; R9: 24 fold higher: *p*<0.001) showing that RN-1 reversed the repression of the γ-globin expression at all stages of erythroid differentiation.

**Fig 2.**
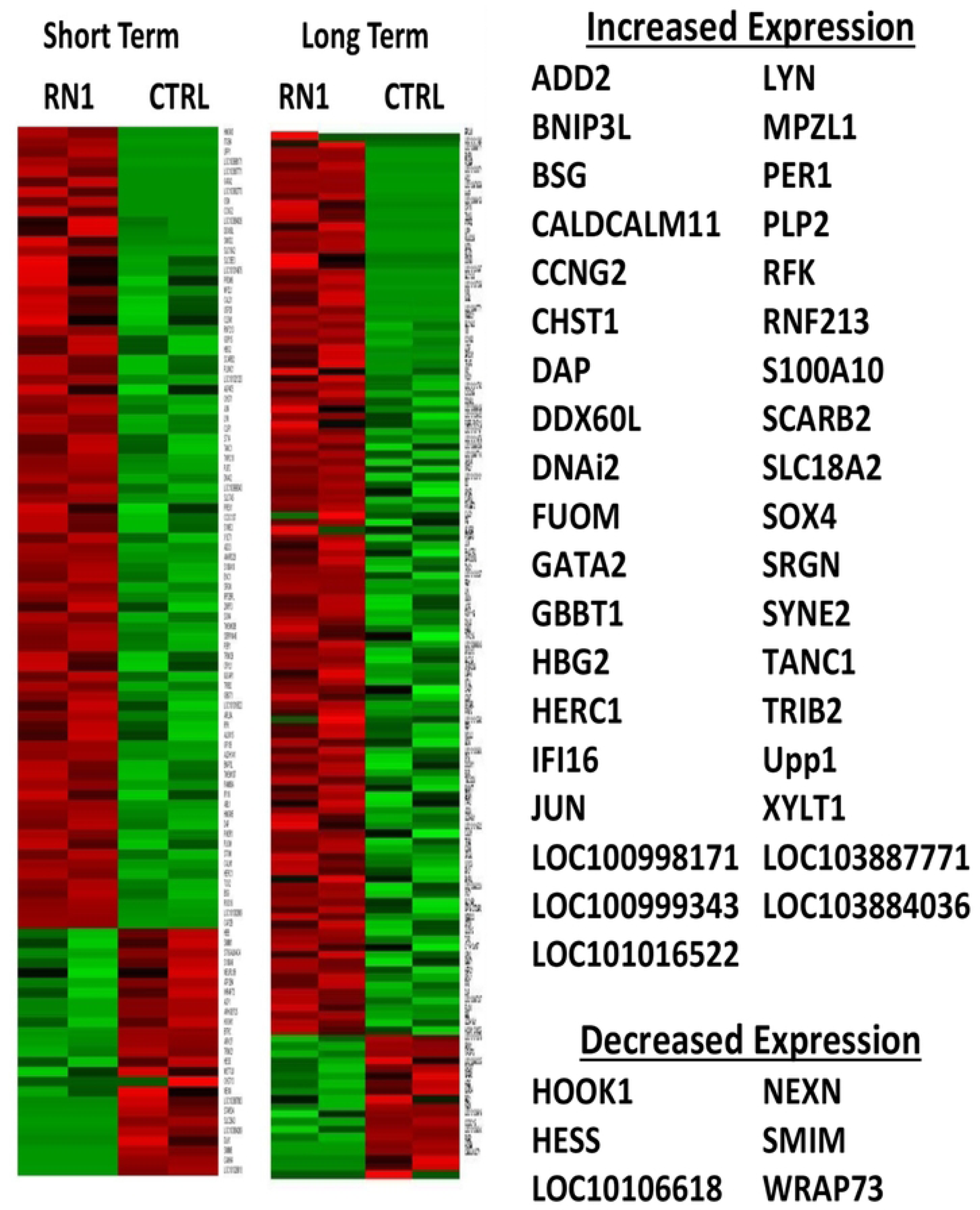
Expression of γ-globin in BM subpopulations. **A.** Expression of γ-globin RNA (γ/γ+β) in subpopulations enriched in BFUe (R6), CFUe (R5), proerythroblasts (R7), and ortho- and polychromatic terminal erythroid cells (R9) isolated from normal untreated baboons (solid fill) and RN-1 treated baboons (diagonal fill). **B.** Fold difference of γ-globin RNA relative to GAPDH RNA (horizontal lines) in isolated BM erythroid cell subpopulations.

### Effect of RN-1 on global gene expression

The effect of RN-1 on global gene expression was investigated by RNAseq analysis of CD34-CD117+bRBC+ FACS-purified CD105+ BM cells (proerythroblasts) of untreated control baboons (n=2) and baboons treated with both short-term (0.25mg/kg/d; 3d; n=2) and long-term schedules (0.25mg/kg/d; 5d/wk; >265d; n=2). The untreated control and short-term treatment samples were obtained from the same two individual baboons, PA897 and PA8548. The pattern of gene expression between these two baboons differed in both the untreated controls (CTRL1, PA8697; CTRL2, PA8548) and short-term RN-1 treatment samples (STT1, PA8697; STT2; PA8548) as revealed by Principal Component Analysis (S2 Fig). This was associated with a 4-5 fold difference in baseline HbF (PA8697, 5.27±0.18% HbF; PA8548, 1.23±0.36% HbF) between the two animals. Gene expression changes due to short term RN-1 treatment were examined by ANOVA analysis to control for differences between the two animals. The long-term treatment samples were obtained from two different baboons (PA8695, LTT1, PA8698 LTT2) that were more related to PA8548 (STT2) than PA 8697 by PCA analysis (S2 Fig) and baseline HbF (PA 8695, 1.10±0.17% HbF; PA8698, 1.02±0.31% HbF). Gene expression differences between these long-term treatment samples were compared directly with the two control samples from PA8697 (CTRL 1) and PA8548 (CTRL 2). In baboons treated for 3d with RN-1 a set of 81 genes was identified whose expression was increased and 24 genes that showed decreased expression compared to the untreated control baboons (S3 Table). In the baboons treated >265d the expression of 119 genes was increased and 18 decreased compared to the untreated controls (S4 Table). Comparison of the gene sets from the short- and long-term treatment regimens identified a common set of 41 genes with increased expression in both treatment regimens while decreased expression of 6 genes was observed (Fig 3). RN-1 treatment therefore affected the expression of a relatively small number of genes, with increased expression observed for most of the genes exhibited changes. Significantly, γ-globin was included among the subset of genes with increased expression in both short and long term treatment regimens. Genes with known roles in erythroid differentiation increased with RN-1 treatment included GFi1b, GATA2, and LYN. RT-PCR assays validated that the expression of DAP, ABL, Gfi-1b, LYN and GATA2 were significantly increased in the BM CD117+bRBC+ subpopulation in RN-1 treated baboons relative to untreated controls. GATA2 exhibited the greatest fold increase in expression (S5 Fig).

**Fig 3.**
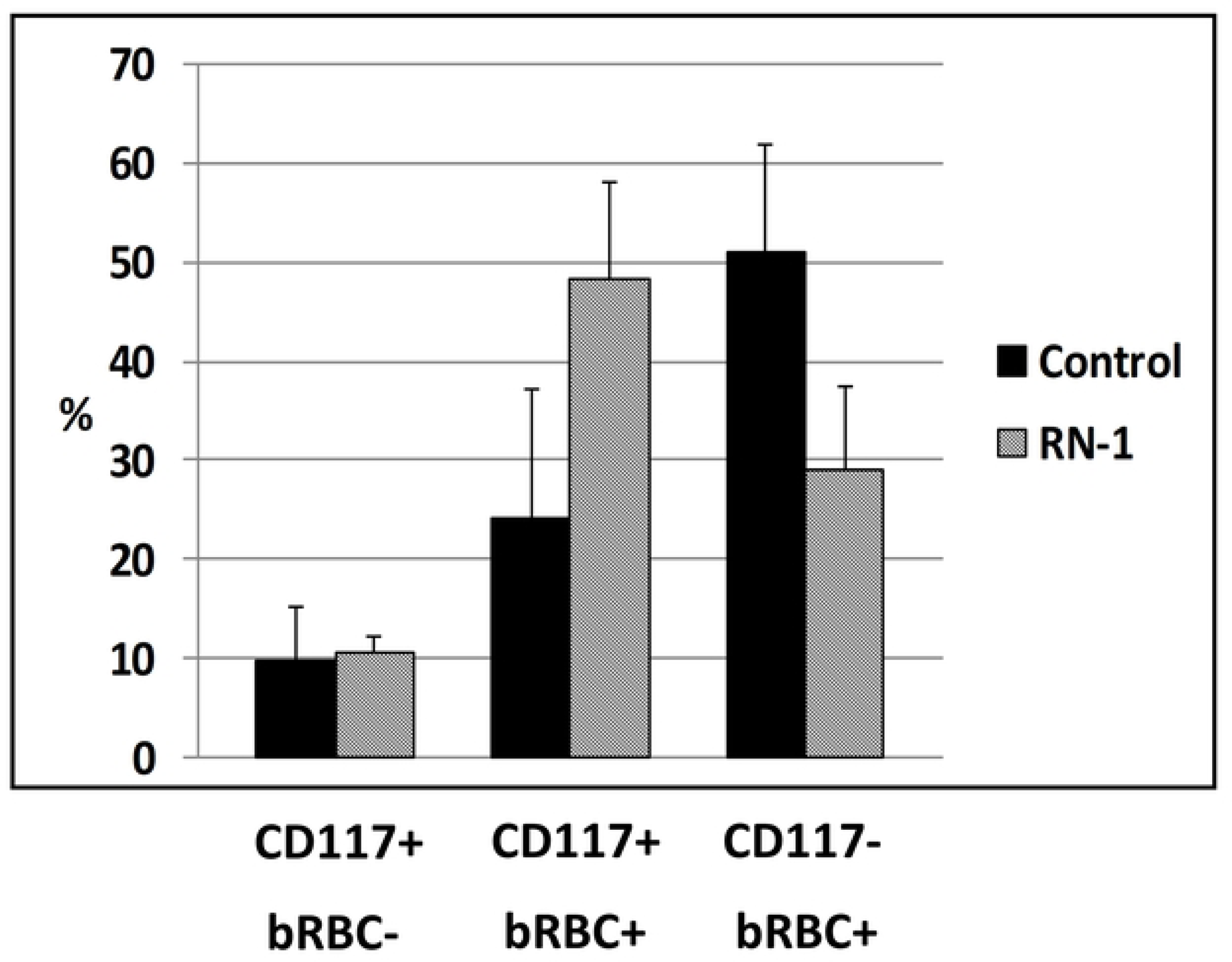
Effect of RN-1 on Gene Expression in Baboon BM Proerythroblasts. Heat map showing differences in gene expression in bone marrow proerythroblast (R7; CD105+CD34-CD117+bRBC+) subpopulations from control, untreated baboons and from baboons treated with either a short term 3d (n=2) or a long term course (n=2) of RN-1 compared to untreated normal control baboons.

### Effect of RN-1 on erythroid differentiation

Additional experiments were performed to determine to whether RN-1 treatment was associated with changes in erythroid differentiation. The effect of the LSD1 inhibitor RN-1 on erythroid differentiation was measured by flow cytometry in baboon BM aspirates following RN-1 administration. Samples from six untreated control baboons were compared to four baboons treated with RN-1. RN-1 treatment (0.25mg/kg/d; 3d) increased the proportion of CD105+CD117+bRBC+BM cells (proerythroblasts) (24.2% pre-treatment; 48.4% post-treatment; p<0.02) and decreased the proportion of CD105+CD117-bRBC+ terminal erythroid cells (51.1% pre-treatment; 29.1% post-treatment; p<0.02, Figure 4B).

**Fig 4.**
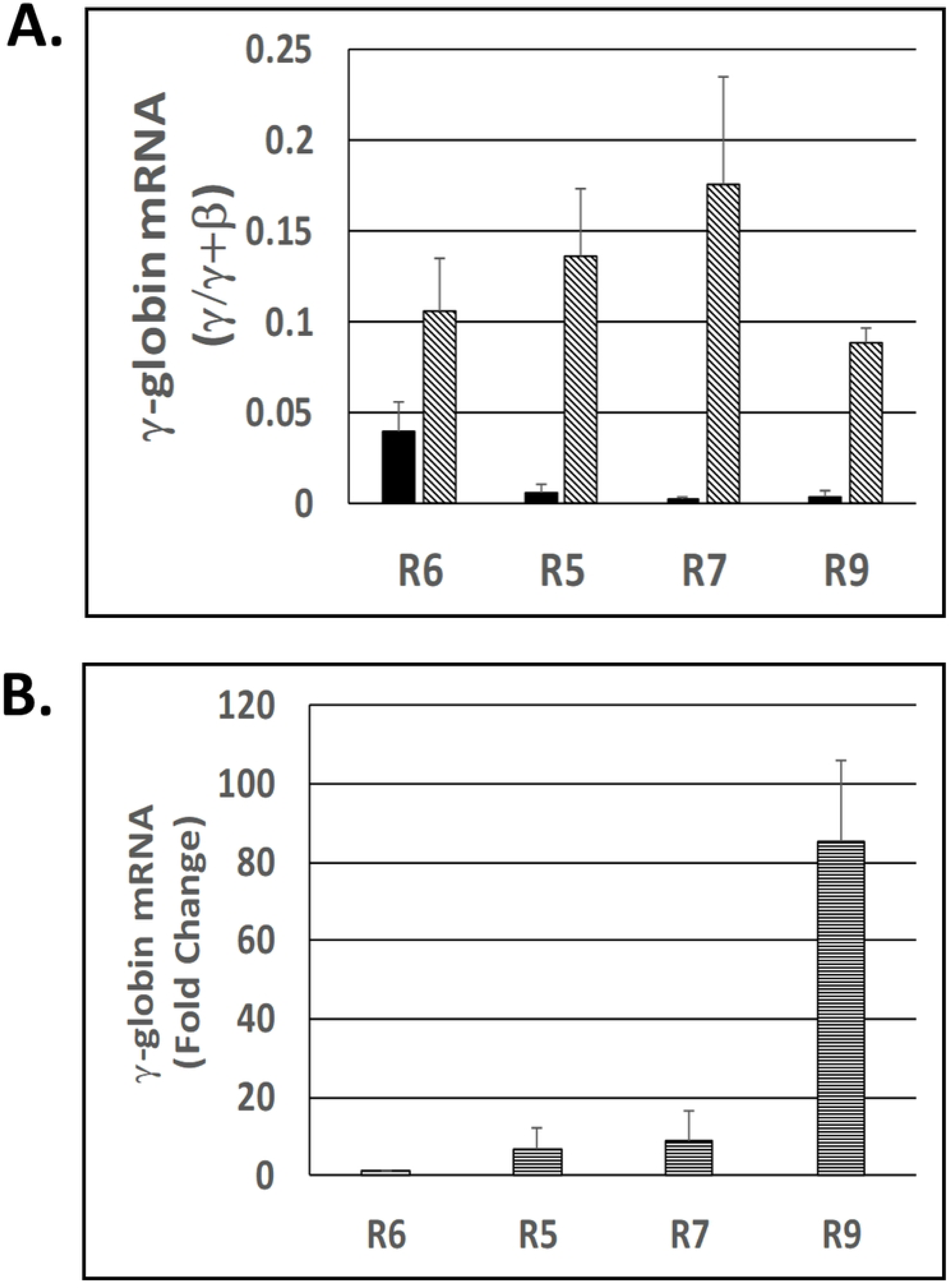
Effect of RN-1 on Erythroid Differentiation. Flow cytometric assay showing differences in distribution of CD117+bRBC-, CD117+bRBC+ and CD117-bRBC+ erythroid cells in bone marrow aspirates from normal, untreated control baboons (filed bars) and RN-1 treated baboons (diagonal bars; mean±SD).

## Discussion

A purification scheme based on immunomagnetic cell separation and FACS was developed to isolate subpopulations of BM erythroid cells enriched for different stages of erythroid differentiation to investigate the mechanism of action of RN-1. The effect of RN1 treatment of experimental baboons on γ-globin expression within each subpopulation was determined. Our results, consistent with previous studies [1,2], show that γ-globin is expressed at higher levels relative to β-globin in early BFUe-enriched progenitors compared to later stages of erythroid differentiation, suggesting that γ-globin repression is initiated after the BFUe stage and is largely completed by the time cells reach the CFUe stage. RN-1 treatment of baboons interferes with this repressive event and allows increased γ-globin relative to β-globin expression throughout all stages of erythroid differentiation in BM cells directly isolated from RN-1 baboons.

Our previous work showed that administration of the LSD1 inhibitor RN-1 to baboons caused large increases in γ-globin expression [20,21]. LSD1 inhibitors were initially tested for their ability to increase HbF in baboons based on evidence that LSD1 was a component of a multi-protein complex is recruited to the γ-globin promoter by the TR2/TR4 orphan nuclear receptor proteins and BCL11A [10,11]. The effect of LSD1 depletion by shRNA and inhibition of its demethylase activity by treatment with LSD1 inhibitors to increase γ-globin expression in primary human CD34+ erythroid cell cultures provided the initial evidence that LSD1 was a potential therapeutic target [15]. In that report and data from our in vivo baboon experiments [20] showed that increased γ-globin expression was accompanied by the acquisition of activating histone methylation marks from the Histone H3 K4 residue at the γ-globin promoter suggesting that the mechanism of LSD1-mediated γ-globin repression was likely due to its ability to remove these activating marks. Modification of the histone methylation status at the γ-globin promoter is associated and may be necessary for increased expression but may not be alone sufficient. Recent experiments in the HuDep-2 cell line where BCL11A was depleted by Protac-mediated targeted degradation suggested that DNA methylation is not sufficient to repress γ-globin expression, [44] but these results could also be the result of the inability of the NURD complex containing MBD2, a “reader” of DNA methylation required for γ-globin repression, to be recruited recruited to the γ-globin promoter in the absence of BCL11A [45].

Our analysis of the effect of in vivo RN-1 treatment on the transcriptome of an isolated proerythroblast BM subpopulation clearly shows that RN-1 increases the expression of a relatively small number of genes, with only few genes exhibiting downregulation. Downregulation of the known γ-globin trans-acting repressors including BCL11A, ZBTB7A, and TR2/TR4 was not observed suggesting that the mechanism responsible for increased γ-globin expression by RN-1 does not involve decreased transcriptional expression of these genes. RN-1 increased PGC-1α in SCD mice [17] while decreased expression of NCOR1 and SOX6 was observed in erythroid cultures derived from β^O^-thalassemia/HbE patients [46]. It is possible that discrepancies between these studies and our own may be due to species differences, culture conditions or dosing.

Recent studies have demonstrated major differences in the abundance of transcription factors, co-activators, and co-repressors between the RNA and protein levels [47, 48]. The targets of LSD1-mediated demethylation are not limited to histone proteins. [49] Lysine methylation can target proteins for proteolytic degradation [50]. It has been observed that LSD1 decreases methylation of DNMT1 and increases its abundance while LSD1 inhibitors reverse this process [51]. The DNMT1 inhibitor decitabine, another powerful HBF-inducing drug, also causes in proteolysis and depletion of the DNMT1 protein [52]. DNMT1 can act as a scaffold protein for multi-protein corepressor complexes, suggesting that depletion of DNMT1 by either decitabine or LSD1 could alter the integrity and/or abundance of the co-repressor complex and thus the recruitment of other epigenome-modifying enzymes and/or readers to the γ-globin promoter [53]. Further knowledge of the effect of LSD1 inhibitors on the proteome are therefore required for a more complete understanding of the mechanism of action of these drugs.

Many of the genes whose transcriptional expression is increased following RN-1 treatment play mechanistic roles in erythroid differentiation, including GATA2, Gfi1B, and LYN. The ability of RN-1 to increase their expression may be responsible, at least in part, for inhibition of erythroid differentiation we observe in the BM of RN-1 treated baboons. LSD1 has been identified as a key factor in repression of the GATA2 gene. LSD1-knockdown increases GATA2 and suppresses GATA1 expression to inhibit erythroid differentiation [54,55]. Gfi-1b drives erythroid differentiation by repression of target genes in a process requiring LSD1 [56]. The increased Gf1-1B expression observed in RN-1 treated BM cells may be due to interference with its autoregulation [57]. LYN, another gene exhibiting increased expression in RN-1 treated baboons, also affects erythroid differentiation. Red cell differentiation is abnormal in mice deficient in LYN [58] and erythroid differentiation is delayed in gain-of-function LYN mice [59]. Therefore, the ability of LSD1 inhibitors to perturb erythroid differentiation is associated with effects on multiple genes.

## Acknowledgements

Library construction and RNA-seq analysis was performed by the UIC Genome Research Core.

